## Supplementary material for "Loss function of tumor suppressor FRMD8 confers resistance to tamoxifen therapy via a dual mechanism": Supplymental information

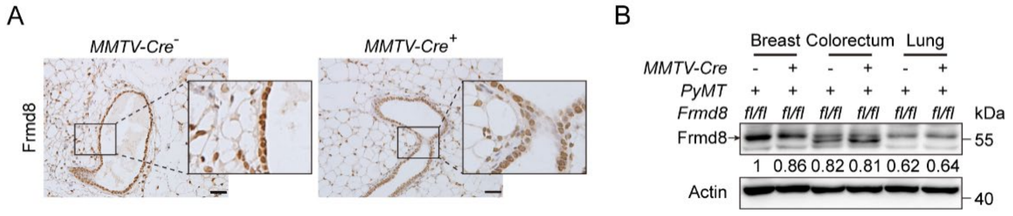


**Figure S1. Loss of FRMD8 promotes breast cancer cell proliferation**

(A) Immunohistochemistry staining for Frmd8 expression in mammary glands from *PyMT* mice. The black boxes represent the magnified typical staining of the original images. Scale bar, 50 μm. (B) Lysates from different tissues of *MMTV-Cre*$^-$; *Frmd8*$^{fl/fl}$; *PyMT* and *MMTV-Cre*$^+$; *Frmd8*$^{fl/fl}$; *PyMT* mice were examined by Western blot. Specific bands are marked with an arrow.

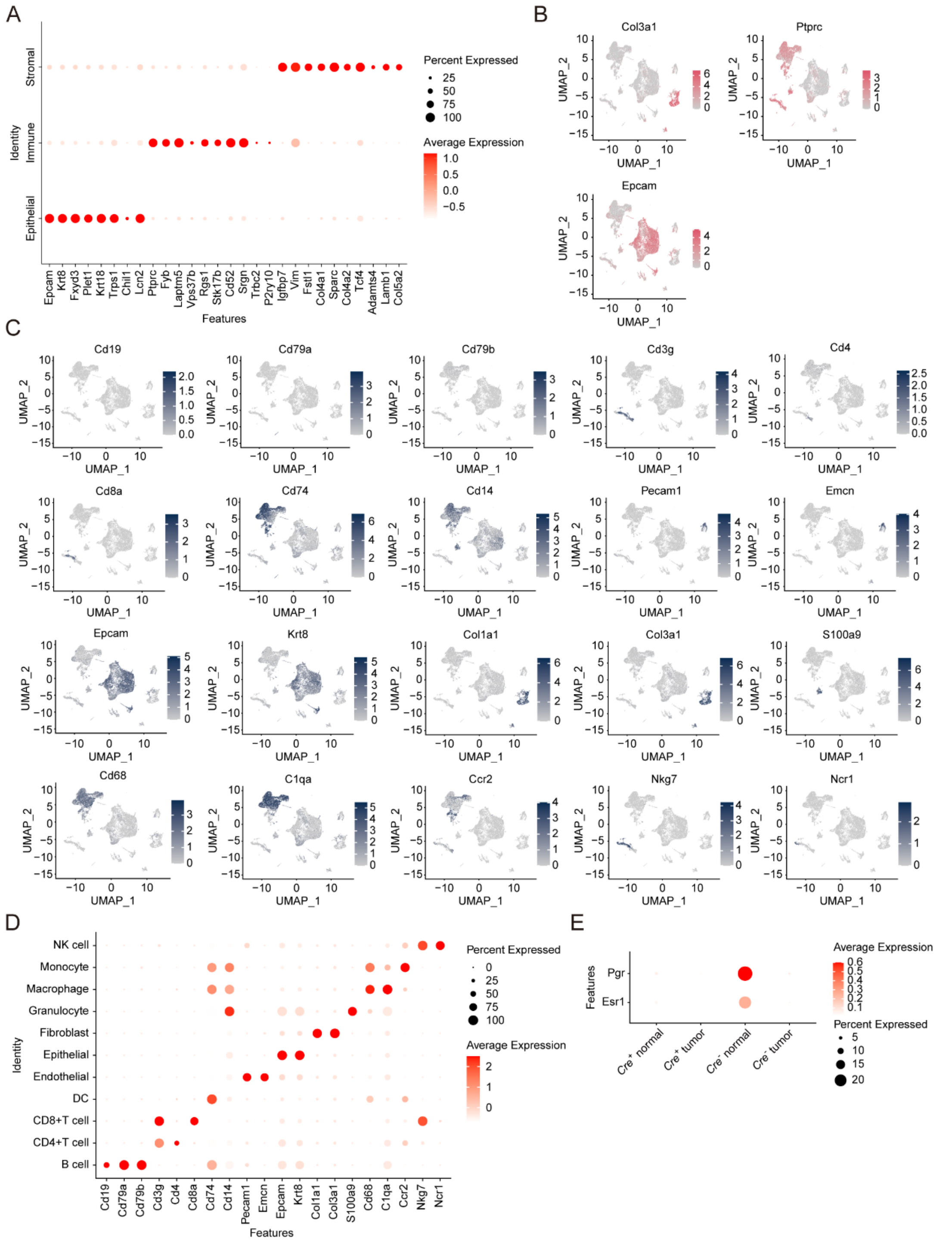


**Figure S2. Distinct cell lineages determined by single-cell RNA-Seq analysis**

(A) Dot plot representing the expression level (red jet) and the number of expressing cells (dot size) of the top markers from main cell lineages. (B) T-SNE visualization of the expression level of main lineages markers. The color key from gray to pink indicates low to high gene expression. (C) T-SNE visualization of the expression level of the top markers from each of the cell lineages. The color key from gray to blue indicates low to high gene expression. (D) Dot plot representing the expression level (red jet) and the number of expressing cells (dot size) of the top markers from each of the cell lineages. (E) Dot plot showing the expression of *Esr1* and *Pgr* in normal and tumor cells from *MMTV-Cre*$^{-}$; *Frmd8*$^{fl/fl}$; *PyMT* and *MMTV-Cre*$^{+}$; *Frmd8*$^{fl/fl}$; *PyMT* mice.

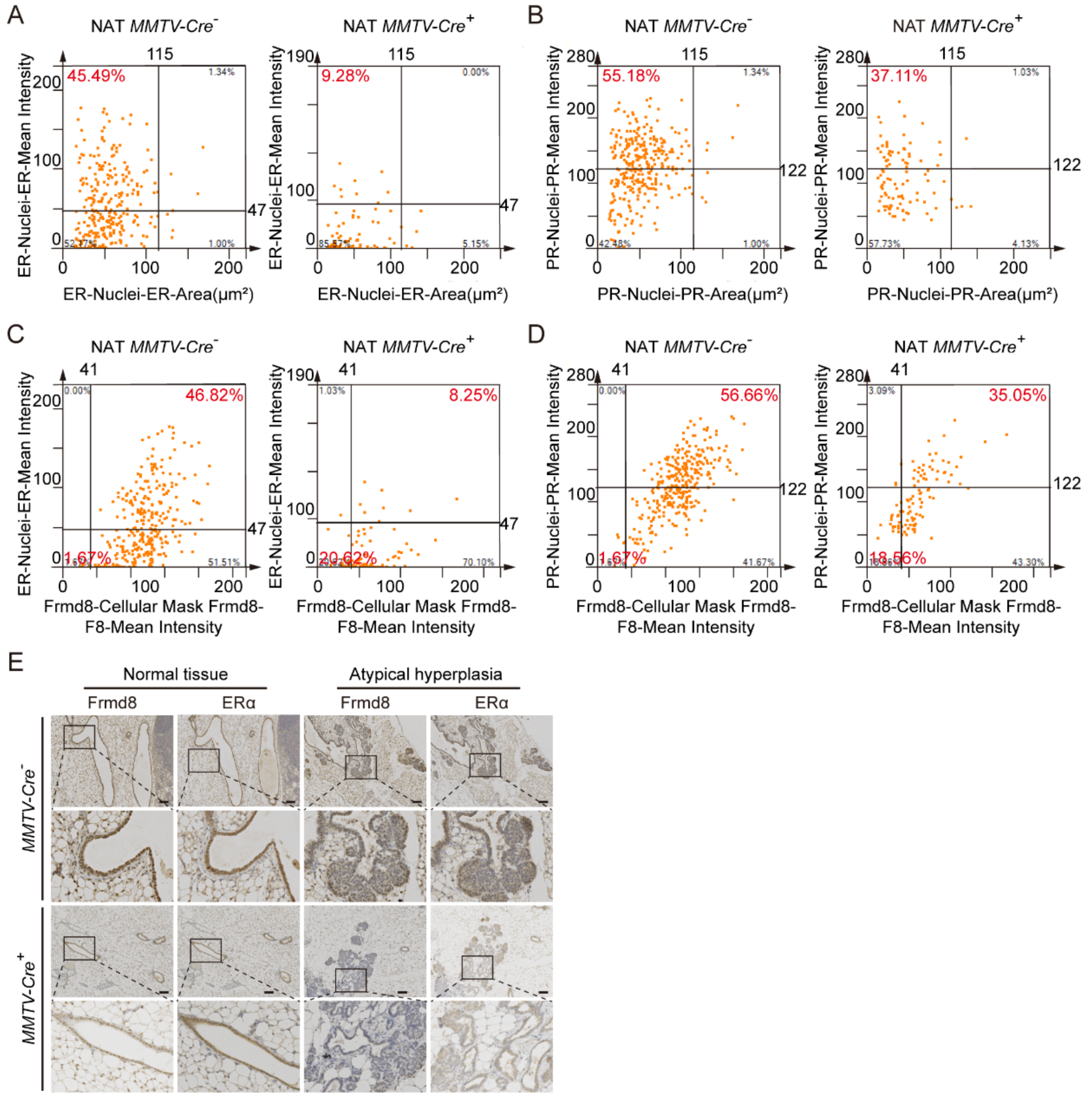


**Figure S3. The expressions of ERα and PR decreased in Frmd8-deleted mammary tissues**

(A-D) The scatter plot shows the cells in multiplex immunofluorescence images of normal tissues adjacent to tumor in *MMTV-Cre*⁻; *Frmd8*$^{fl/fl}$; *PyMT* and *MMTV-Cre*⁺; *Frmd8*$^{fl/fl}$; *PyMT* mice. (E) IHC staining for Frmd8 and ERα expression in 7-week-old mammary glands from *PyMT* mice. The black boxes represent the magnified typical staining of the original images. Scale bar, 100 μm.

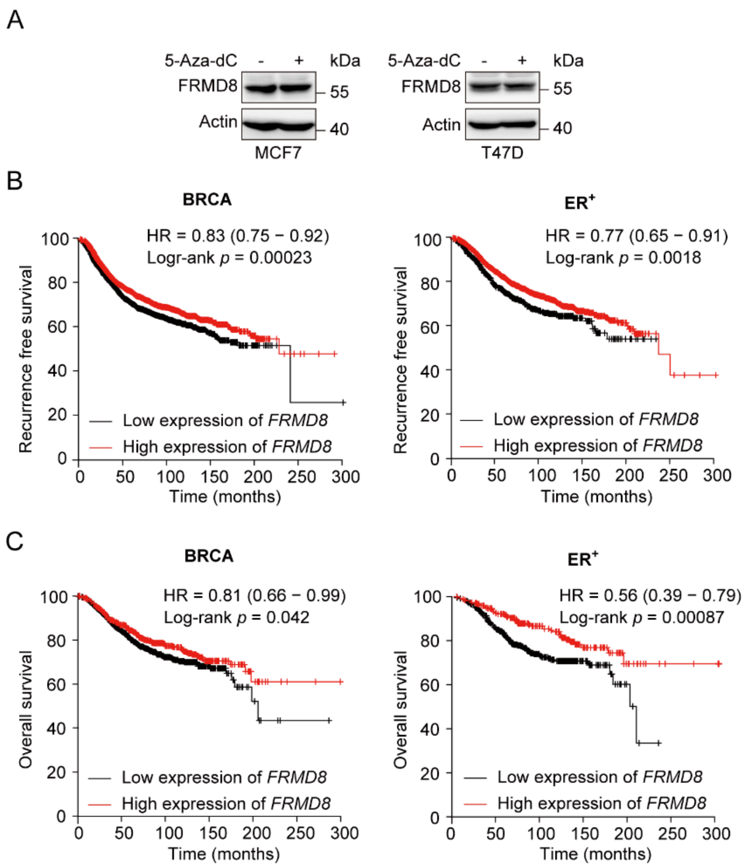


**Figure S4. *FRMD8* promoter is methylated and low FRMD8 level predicts poor prognosis in breast cancer patients**

(A) MCF7 and T47D cells were treated with 5-Aza-dC (10 μM) for 48 h. Protein expression of FRMD8 was examined by Western blot. (B-C) Recurrence free survival (B) and overall survival (C) of breast cancer patients with tamoxifen treatment according to *FRMD8* expression were analyzed according to Kaplan-Meier plotter.

Table S2. Primers for genotyping, RNA silencing and qRT-PCR

| **Genotyping** | | |
|---|---|---|
| **Mice** | **Forward** | **Reverse** |
| *Frmd8* floxed mice | 5’-TGGGTGGAGGTGACAAGGCAAGAA-3’ | 5’-CAGGCTCAGGGTGGTTAGGTCAAT-3’ |
| *MMTV-Cre* transgenic mice | 5’-ATTTGCCTGCATTACCGGTCG-3’ | 5’-CAGCATTGCTGTCACTTGGTC-3’ |
| *MMTV-PyMT* transgenic mice | 5’-GGAAGCAAGTACTTCACAAGGG-3’ | 5’-GGAAAGTCACTAGGAGCAGGG-3’ |
| **siRNA targeted sequence** | | |
| siFRMD8#1 | CCAAGCAGGCCGAACTGAT | |
| siFRMD8#2 | TGCTCTATGAGGAGGCCAA | |
| siFOXO3A | CAGCGGAGCTCTAGCTTCCCGTATA | |
| siUBE3A | CCTACATCTCATACTTGCTTT | |
| **qRT-PCR** | | |
| **Gene** | **Forward** | **Reverse** |
| Mouse *Frmd8* | 5’-CGATGATGATGTCGCCATGG-3’ | 5’-AGTCCTCCAAGTCACAAGGG-3’ |
| Mouse *Gapdh* | 5’-GGGTCCCAGCTTAGGTTCAT-3’ | 5’-CATTCTCGGCCTTGACTGTG-3’ |
| Human *FRMD8* | 5’-TTCTTCCACGGTGAGGTTGA-3’ | 5’-TCGAACTCCAGCCACAAGAT-3’ |
| Human *ESR1* | 5’-ATGTGCCTGGCTAGAGATCC-3’ | 5’-CAAACTCCTCTCCCTGCAGA-3’ |
| Human *GAPDH* | 5’-GCACCACCAACTGCTTAGCA-3’ | 5’-TCTTCTGGGTGGCAGTGATG-3’ |
| **Chip-qPCR** | | |
| **Gene** | **Forward** | **Reverse** |
| Human *ESR1* | 5’-CCACTGGGAAATGAGAGACCTCGT-3’ | 5’-GTGGATCAAATGCCTTACTGGCCC-3’ |

Table S3. Patient information of tissue microarray

| Tissue code | Gender | Age | Tumor location | Pathologic classification | Primary or metastasis | TNM category | | | AJCC stage | Pathologic stage | State of survival | Survival time (month) | Expression of FRMD8 | Expression of ERα |
|---|---|---|---|---|---|---|---|---|---|---|---|---|---|---|
| J07A0929 | Female | 48 | Breast | Invasive ductal carcinoma | Primary | T2 | N0 | M0 | 2A | Ⅱ | Alive | 150 | 2 | Positive |
| J07A0930 | Female | 60 | Breast | Invasive ductal carcinoma | Primary | T2 | N0 | M0 | 2A | Ⅱ | Alive | 150 | 2 | Positive |
| J07A0933 | Female | 47 | Breast | Invasive ductal carcinoma | Primary | T3 | N2 | M0 | 3A | Ⅰ- Ⅱ | Dead | 32 | 1 | Positive |
| J07A0934 | Female | 37 | Breast | Invasive ductal carcinoma | Primary | T2 | N0 | M0 | 2A | Ⅱ | Alive | 149 | 2 | Positive |
| J07A0935 | Female | 54 | Breast | Invasive ductal carcinoma | Primary | T3 | N1 | M0 | 3A | Ⅱ | Alive | 148 | 0 | Negative |
| J07A0936 | Female | 72 | Breast | Invasive ductal carcinoma | Primary | T2 | N0 | M0 | 2A | Ⅰ- Ⅱ | Alive | 148 | 1 | Positive |
| J07A0937 | Female | 31 | Breast | Invasive ductal carcinoma with Invasive lobular carcinoma | Primary | T2 | N0 | M0 | 2A | Ⅰ- Ⅱ | Dead | 147 | 1 | Positive |
| J07A0939 | Female | 50 | Breast | Invasive ductal carcinoma | Primary | T3 | N3 | M0 | 3C | Ⅱ | Alive | 147 | 3 | Negative |
| J07A0940 | Female | 57 | Breast | Invasive ductal carcinoma | Primary | T2 | N2 | M0 | 3A | Ⅰ- Ⅱ | Alive | 147 | 2 | Positive |
| J07A0941 | Female | 52 | Breast | Invasive ductal carcinoma with Invasive lobular carcinoma and apocrine carcinoma | Primary | T2 | N1 | M0 | 2B | Ⅰ | Alive | 147 | 1 | Positive |
| J07A0942 | Female | 76 | Breast | Invasive ductal carcinoma | Primary | T2 | N0 | M0 | 2A | Ⅱ | Dead | 33 | 1 | Negative |
| J07A0943 | Female | 63 | Breast | Invasive ductal carcinoma | Primary | T1 | N0 | M0 | 1 | Ⅱ | Alive | 146 | 1 | Positive |
| J07A0945 | Female | 82 | Breast | Invasive ductal carcinoma | Primary | T2 | N0 | M0 | 2A | Ⅱ | Dead | 54 | 1 | Negative |
| J07A0946 | Female | 48 | Breast | Invasive ductal carcinoma | Primary | T3 | N2 | M0 | 3A | Ⅰ | Alive | 145 | 1 | Negative |
| J07A0947 | Female | 37 | Breast | Invasive ductal carcinoma | Primary | T2 | N0 | M0 | 2A | Ⅱ-Ⅲ | Dead | 93 | 1 | Weakly positive |
| J07A0951 | Female | 34 | Breast | Invasive ductal carcinoma | Primary | T1 | N0 | M0 | 1 | Ⅱ | Alive | 145 | 2 | Positive |
| J07A0952 | Female | 73 | Breast | Invasive ductal carcinoma with invasive lobular carcinoma | Primary | T2 | N2 | M0 | 3A | Ⅱ-Ⅲ | Dead | 114 | 1 | Positive |

| | | | | | | | | | | | | | | |
|---|---|---|---|---|---|---|---|---|---|---|---|---|---|---|
| J07A0956 | Female | 54 | Breast | Invasive ductal carcinoma | Primary | T2 | N0 | M0 | 2A | II | Dead | 97 | 1 | Negative |
| J07A0957 | Female | 56 | Breast | Invasive ductal carcinoma | Primary | T2 | N0 | M0 | 2A | II | Alive | 144 | 2 | Positive |
| J07A0958 | Female | 82 | Breast | Invasive ductal carcinoma | Primary | T1 | - | M0 | - | II | Alive | 143 | 2 | Positive |
| J07A0960 | Female | 72 | Breast | Invasive ductal carcinoma | Primary | T2 | N2 | M0 | 3A | II | Alive | 143 | 2 | Negative |
| J07A0961 | Female | 33 | Breast | Invasive ductal carcinoma | Primary | T2 | N2 | M0 | 3A | II | Dead | 19 | 1 | - |
| J07A0962 | Female | 63 | Breast | Invasive ductal carcinoma | Primary | T2 | N1 | M0 | 2B | I - II | Alive | 143 | 1 | Negative |
| J07A0963 | Female | 51 | Breast | Invasive ductal carcinoma | Primary | T1 | N2 | M0 | 3A | II | Dead | 18 | 1 | Negative |
| J07A0964 | Female | 44 | Breast | Invasive ductal carcinoma | Primary | T3 | N1 | M0 | 3A | II | Dead | 17 | 1 | Negative |
| J07A0965 | Female | 67 | Breast | Invasive ductal carcinoma | Primary | T1 | N0 | M0 | 1 | I - II | Alive | 143 | 1 | Positive |
| J07A0966 | Female | 48 | Breast | Invasive ductal carcinoma | Primary | T2 | N3 | M0 | 3C | II | Dead | 11 | 0 | Negative |
| J07A0968 | Female | 68 | Breast | Invasive ductal carcinoma | Primary | T1 | N1 | M0 | 2A | I | Dead | 82 | 1 | Negative |
| J07A0969 | Female | 52 | Breast | Invasive ductal carcinoma | Primary | T2 | N0 | M0 | 2A | II | Alive | 142 | 2 | Positive |
| J07A0970 | Female | 55 | Breast | Invasive ductal carcinoma | Primary | T2 | N2 | M0 | 3A | II | Alive | 142 | 1 | Positive |
| J07A0973 | Female | 76 | Breast | Invasive ductal carcinoma | Primary | T3 | - | M0 | - | I | Alive | 141 | 1 | Positive |
| J07A0974 | Female | 72 | Breast | Invasive ductal carcinoma | Primary | T1 | N1 | M0 | 2A | I - II | Alive | 141 | 2 | Positive |
| J07A0976 | Female | 44 | Breast | Invasive ductal carcinoma | Primary | T2 | N0 | M0 | 2A | I - II | Alive | 141 | 2 | Positive |
| J07A0977 | Female | 77 | Breast | Invasive ductal carcinoma | Primary | T2 | N2 | M0 | 3A | II | Alive | 141 | 2 | Positive |
| J07A0978 | Female | 60 | Breast | Invasive ductal carcinoma | Primary | T2 | N2 | M0 | 3A | I - II | Alive | 141 | 1 | Positive |
| J07A0980 | Female | 44 | Breast | Invasive ductal carcinoma | Primary | T2 | N1 | M0 | 2B | II | Alive | 140 | 2 | Positive |
| J07A0981 | Female | 38 | Breast | Invasive ductal carcinoma | Primary | T2 | N2 | M0 | 3A | I - II | Alive | 140 | 2 | Positive |
| J07A0983 | Female | 57 | Breast | Invasive ductal carcinoma | Primary | T1 | N1 | M0 | 2A | I - II | Alive | 139 | 2 | Negative |
| J07A0985 | Female | 64 | Breast | Invasive ductal carcinoma | Primary | T2 | N1 | M0 | 2B | II | Alive | 138 | 1 | Positive |
| J07A0986 | Female | 44 | Breast | Invasive ductal carcinoma | Primary | T3 | N2 | M0 | 3A | II | Alive | 138 | 2 | Negative |
| J07A0987 | Female | 48 | Breast | Invasive ductal carcinoma | Primary | T2 | N0 | M0 | 2A | I | Dead | 32 | 0 | Positive |
| J07A0988 | Female | 47 | Breast | Invasive ductal carcinoma | Primary | T2 | N0 | M0 | 2A | I | Alive | 138 | 2 | Positive |

| | | | | | | | | | | | | | | |
|---|---|---|---|---|---|---|---|---|---|---|---|---|---|---|
| J07A0989 | Female | 47 | Breast | Invasive ductal carcinoma with invasive lobular carcinoma | Primary | T1 | N1 | M0 | 2A | Ⅰ | Alive | 137 | 0 | Positive |
| J07A0990 | Female | 33 | Breast | Invasive ductal carcinoma | Primary | T3 | N3 | M0 | 3C | Ⅱ | Alive | 137 | 0 | Positive |
| J07A0991 | Female | 46 | Breast | Invasive ductal carcinoma | Primary | T2 | N2 | M0 | 3A | Ⅱ | Dead | 7 | 1 | Positive |
| J07A0992 | Female | 39 | Breast | Invasive ductal carcinoma | Primary | T2 | N3 | M0 | 3C | Ⅰ | Alive | 137 | 1 | Positive |
| J07A0993 | Female | 46 | Breast | Invasive ductal carcinoma | Primary | T2 | N0 | M0 | 2A | Ⅱ | Alive | 137 | 3 | Negative |
| J07A0994 | Female | 40 | Breast | Invasive ductal carcinoma | Primary | T1 | N2 | M0 | 3A | Ⅱ | Alive | 137 | 1 | Positive |
| J07A0997 | Female | 61 | Breast | Invasive ductal carcinoma | Primary | T2 | N0 | M0 | 2A | Ⅱ | Alive | 136 | 1 | Positive |
| J07A0998 | Female | 61 | Breast | Invasive ductal carcinoma | Primary | T1 | N1 | M0 | 2A | Ⅰ | Alive | 136 | 2 | Positive |
| J07A0999 | Female | 48 | Breast | Invasive ductal carcinoma | Primary | T2 | N2 | M0 | 3A | Ⅱ | Dead | 125 | 1 | - |
| J07A1000 | Female | 40 | Breast | Invasive ductal carcinoma | Primary | T2 | N1 | M0 | 2B | Ⅰ | Dead | 23 | 1 | Negative |
| J07A1001 | Female | 62 | Breast | Invasive ductal carcinoma | Primary | T2 | N1 | M0 | 2B | Ⅰ - Ⅱ | Alive | 135 | 1 | Negative |
| J07A1003 | Female | 36 | Breast | Invasive ductal carcinoma | Primary | T1 | N2 | M0 | 3A | Ⅱ | Alive | 135 | 3 | Positive |
| J07A1004 | Female | 38 | Breast | Invasive ductal carcinoma | Primary | T3 | N1 | M0 | 3A | Ⅰ - Ⅱ | Alive | 135 | 2 | Positive |
| J07A1006 | Female | 73 | Breast | Invasive ductal carcinoma | Primary | T2 | N1 | M0 | 2B | Ⅰ - Ⅱ | Alive | 135 | 2 | Negative |
| J07A1007 | Female | 45 | Breast | Invasive ductal carcinoma | Primary | T1 | N1 | M0 | 2A | Ⅰ - Ⅱ | Dead | 47 | 1 | Negative |
| J07A1010 | Female | 76 | Breast | Invasive ductal carcinoma | Primary | T2 | N0 | M0 | 2A | Ⅱ | Dead | 77 | 3 | Negative |
| J07A1011 | Female | 47 | Breast | Invasive ductal carcinoma | Primary | T2 | N1 | M0 | 2B | Ⅱ | Alive | 134 | 2 | Positive |
| J07A1012 | Female | 61 | Breast | Invasive ductal carcinoma | Primary | T2 | N1 | M0 | 2B | Ⅱ | Alive | 134 | 1 | Positive |
| J07A1013 | Female | 78 | Breast | Invasive ductal carcinoma | Primary | T2 | N0 | M0 | 2A | Ⅱ | Dead | 92 | 2 | Positive |
| J07A1014 | Female | 55 | Breast | Invasive ductal carcinoma | Primary | T2 | N1 | M0 | 2B | Ⅰ | Dead | 31 | 2 | Positive |
| J07A1015 | Female | 50 | Breast | Invasive ductal carcinoma | Primary | T2 | N3 | M0 | 3C | Ⅱ | Dead | 78 | 1 | - |
| J07A1017 | Female | 66 | Breast | Invasive ductal carcinoma | Primary | T2 | N0 | M0 | 2A | Ⅱ | Alive | 133 | 2 | Positive |
| J07A1019 | Female | 48 | Breast | Invasive ductal carcinoma | Primary | T2 | N0 | M0 | 2A | Ⅰ - Ⅱ | Alive | 132 | 2 | Positive |

| | | | | | | | | | | | | | | |
|---|---|---|---|---|---|---|---|---|---|---|---|---|---|---|
| J07A1020 | Female | 41 | Breast | Invasive ductal carcinoma with invasive lobular carcinoma | Primary | T1 | N0 | M0 | 1 | II | Alive | 132 | 2 | Positive |
| J07A1021 | Female | 51 | Breast | Invasive ductal carcinoma | Primary | T2 | N0 | M0 | 2A | II | Alive | 132 | 1 | Positive |
| J07A1022 | Female | 45 | Breast | Invasive ductal carcinoma | Primary | T2 | N0 | M0 | 2A | II | Alive | 132 | 2 | Positive |
| J07A1023 | Female | 75 | Breast | Invasive ductal carcinoma | Primary | T2 | N2 | M0 | 3A | II -III | Dead | 53 | 2 | Positive |
| J07A1024 | Female | 47 | Breast | Invasive ductal carcinoma | Primary | T2 | N0 | M0 | 2A | II | Alive | 131 | 2 | Positive |
| J07A1026 | Female | 56 | Breast | Invasive ductal carcinoma | Primary | T2 | N1 | M0 | 2B | II | Alive | 131 | 1 | Positive |
| J07A1028 | Female | 74 | Breast | Invasive ductal carcinoma | Primary | T3 | N2 | M0 | 3A | I - II | Dead | 23 | 1 | Positive |
| J07A1030 | Female | 50 | Breast | Invasive ductal carcinoma | Primary | T1 | N2 | M0 | 3A | I - II | Alive | 130 | 2 | Positive |
| J07A1031 | Female | 72 | Breast | Invasive ductal carcinoma | Primary | T2 | N1 | M0 | 2B | II | Alive | 130 | 2 | Positive |
| J07A1032 | Female | 36 | Breast | Invasive ductal carcinoma | Primary | T2 | N3 | M0 | 3C | II | Dead | 15 | 3 | Negative |
| J07A1033 | Female | 46 | Breast | Invasive ductal carcinoma | Primary | T2 | N1 | M0 | 2B | I | Alive | 129 | 2 | Positive |
| J07A1034 | Female | 54 | Breast | Invasive ductal carcinoma | Primary | T2 | N0 | M0 | 2A | II | Dead | 60 | 3 | Positive |
| J07A1036 | Female | 63 | Breast | Invasive ductal carcinoma | Primary | T2 | N1 | M0 | 2B | II -III | Dead | 110 | 1 | Positive |
| J07A1038 | Female | 50 | Breast | Invasive ductal carcinoma | Primary | T2 | N1 | M0 | 2B | II | Alive | 128 | 3 | Positive |
| J07A1040 | Female | 53 | Breast | Invasive ductal carcinoma | Primary | T2 | N2 | M0 | 3A | II | Dead | 63 | 2 | Positive |
| J07A1041 | Female | 63 | Breast | Invasive ductal carcinoma | Primary | T1 | N1 | M0 | 2A | II | Dead | 35 | 2 | Negative |
| J07A1043 | Female | 37 | Breast | Invasive ductal carcinoma | Primary | T1 | N3 | M0 | 3C | II | Alive | 128 | 3 | Positive |
| J07A1045 | Female | 55 | Breast | Invasive ductal carcinoma | Primary | T2 | N1 | M0 | 2B | II | Alive | 128 | 2 | Positive |
| J07A1046 | Female | 40 | Breast | Invasive ductal carcinoma | Primary | T2 | N0 | M0 | 2A | I - II | Dead | 44 | 2 | Positive |
| J07A1047 | Female | 29 | Breast | Invasive ductal carcinoma | Primary | T2 | N2 | M0 | 3A | II | Dead | 78 | 1 | Negative |
| J07A1052 | Female | 45 | Breast | Invasive ductal carcinoma | Primary | T2 | N1 | M0 | 2B | II | Alive | 127 | 1 | Positive |
| J07A1053 | Female | 31 | Breast | Invasive ductal carcinoma | Primary | T3 | N2 | M0 | 3A | II | Dead | 4 | 2 | Negative |
| J07A1055 | Female | 54 | Breast | Invasive ductal carcinoma | Primary | T2 | N2 | M0 | 3A | III | Dead | 4 | 2 | Positive |
| J07A1057 | Female | 48 | Breast | Invasive ductal carcinoma | Primary | T2 | N1 | M0 | 2B | II | Alive | 126 | 2 | Positive |

| | | | | | | | | | | | | | | |
|---|---|---|---|---|---|---|---|---|---|---|---|---|---|---|
| J07A1059 | Female | 71 | Breast | Invasive ductal carcinoma | Primary | T2 | N0 | M0 | 2A | II | Dead | 62 | 2 | Negative |
| J07A1060 | Female | 49 | Breast | Invasive ductal carcinoma | Primary | T2 | N0 | M0 | 2A | II | Alive | 120 | 2 | Negative |
| J07A1061 | Female | 69 | Breast | Invasive ductal carcinoma | Primary | T2 | N1 | M0 | 2B | II | Alive | 120 | 2 | Positive |
| J07A1062 | Female | 46 | Breast | Invasive ductal carcinoma | Primary | T1 | N1 | M0 | 2A | II | Alive | 119 | 3 | Negative |
| J07A1065 | Female | 48 | Breast | Invasive ductal carcinoma | Primary | T2 | N1 | M0 | 2B | II -III | Dead | 19 | 1 | - |
| J07A1067 | Female | 82 | Breast | Invasive ductal carcinoma | Primary | T3 | N2 | M0 | 3A | II | Dead | 2 | 1 | Negative |
| J07A1068 | Female | 52 | Breast | Invasive ductal carcinoma | Primary | T2 | N2 | M0 | 3A | II | Dead | 110 | 2 | Negative |
| J07A1070 | Female | 44 | Breast | Invasive ductal carcinoma | Primary | T2 | N2 | M0 | 3A | II | Alive | 118 | 1 | Positive |
| J07A1073 | Female | 51 | Breast | Invasive ductal carcinoma | Primary | T2 | N0 | M0 | 2A | II | Alive | 118 | 2 | Positive |
| J07A1074 | Female | 43 | Breast | Invasive ductal carcinoma | Primary | T3 | N2 | M0 | 3A | II | Alive | 117 | 1 | Negative |
| J07A1076 | Female | 37 | Breast | Invasive ductal carcinoma | Primary | T1 | N0 | M0 | 1 | II -III | Dead | 85 | 3 | Weakly positive |
| J07A1080 | Female | 65 | Breast | Invasive ductal carcinoma | Primary | T1 | N0 | M0 | 1A | | Alive | 115 | 1 | - |
| J07A1081 | Female | 51 | Breast | Invasive ductal carcinoma with invasive lobular carcinoma | Primary | T2 | N0 | M0 | 2A | I - II | Alive | 115 | 1 | Positive |
| J07A1083 | Female | 40 | Breast | Invasive ductal carcinoma | Primary | T2 | N0 | M0 | 2A | - | Alive | 115 | 0 | - |
| J07A1084 | Female | 64 | Breast | Invasive ductal carcinoma | Primary | T2 | N0 | M0 | 2A | II | Dead | 68 | 2 | Negative |
| J07A1085 | Female | 52 | Breast | Invasive ductal carcinoma | Primary | T2 | N3 | M0 | 3C | II | Dead | 61 | 1 | Positive |
| J07A1086 | Female | 66 | Breast | Invasive ductal carcinoma | Primary | T1 | N1 | M0 | 2A | II -III | Alive | 115 | 2 | - |
| J07A1087 | Female | 68 | Breast | Invasive ductal carcinoma | Primary | T2 | N1 | M0 | 2B | II | Alive | 115 | 0 | Negative |
| J07A1088 | Female | 39 | Breast | Invasive ductal carcinoma | Primary | T2 | N2 | M0 | 3A | II | Alive | 114 | 1 | Negative |
| J07A1090 | Female | 41 | Breast | Invasive ductal carcinoma | Primary | T2 | N1 | M0 | 2B | I - II | Alive | 114 | 2 | Negative |
| J07A1093 | Female | 61 | Breast | Invasive ductal carcinoma | Primary | T2 | N0 | M0 | 2A | II | Alive | 113 | 1 | Positive |
| J07A1095 | Female | 65 | Breast | Invasive ductal carcinoma | Primary | T2 | N0 | M0 | 2A | II | Alive | 112 | 1 | Positive |
| J07A1096 | Female | 60 | Breast | Invasive ductal carcinoma | Primary | T2 | N0 | M0 | 2A | II | Alive | 112 | 1 | Positive |

| | | | | | | | | | | | | | | |
|---|---|---|---|---|---|---|---|---|---|---|---|---|---|---|
| J07A1097 | Female | 50 | Breast | Invasive ductal carcinoma | Primary | T2 | N2 | M0 | 3A | II | Alive | 112 | 3 | Negative |
| J07A1098 | Female | 42 | Breast | Invasive ductal carcinoma | Primary | T2 | N1 | M0 | 2B | II | Alive | 112 | 1 | Negative |
| J07A1099 | Female | 40 | Breast | Invasive ductal carcinoma | Primary | T2 | N0 | M0 | 2A | II | Alive | 112 | 2 | Positive |
| J07A1100 | Female | 40 | Breast | Invasive ductal carcinoma | Primary | T2 | N0 | M0 | 2A | II | Alive | 112 | 2 | Weakly positive |
| J07A1101 | Female | 57 | Breast | Invasive ductal carcinoma | Primary | T1 | N0 | M0 | 1 | I - II | Alive | 111 | 3 | Positive |
| J07A1102 | Female | 47 | Breast | Invasive ductal carcinoma | Primary | T2 | N0 | M0 | 2A | II | Alive | 111 | 0 | Positive |
| J07A1103 | Female | 67 | Breast | Invasive ductal carcinoma | Primary | T2 | N1 | M0 | 2B | II | Alive | 111 | 1 | Positive |
| J07A1106 | Female | 46 | Breast | Invasive ductal carcinoma | Primary | T2 | N2 | M0 | 3A | II | Alive | 111 | 1 | Positive |
| J07A1107 | Female | 34 | Breast | Invasive ductal carcinoma | Primary | T2 | N1 | M0 | 2B | II | Alive | 110 | 2 | Positive |
| J07A1108 | Female | 42 | Breast | Cribriform carcinoma | Primary | T2 | N0 | M0 | 2A | II | Alive | 110 | 1 | - |
| J07A1109 | Female | 73 | Breast | Invasive ductal carcinoma | Primary | T1 | N2 | M0 | 3A | II | Dead | 79 | 1 | Positive |
| J07A1110 | Female | 56 | Breast | Invasive ductal carcinoma with invasive lobular carcinoma | Primary | T2 | N2 | M0 | 3A | II | Dead | 80 | 1 | Positive |
| J07A1111 | Female | 59 | Breast | Invasive ductal carcinoma | Primary | T1 | N0 | M0 | 1 | II | Alive | 110 | 2 | Negative |
| J07A1113 | Female | 53 | Breast | Invasive ductal carcinoma | Primary | T1 | N0 | M0 | 1 | II | Alive | 110 | 1 | Positive |
| J07A1115 | Female | 72 | Breast | Invasive ductal carcinoma | Primary | T1 | N1 | M0 | 2A | II | Dead | 46 | 0 | Negative |
| J07A1117 | Female | 39 | Breast | Mucinous carcinoma | Primary | T2 | N0 | M0 | 2A | - | Alive | 109 | 2 | - |
| J07A1118 | Female | 52 | Breast | Invasive ductal carcinoma | Primary | T2 | N1 | M0 | 2B | II | Alive | 109 | 1 | Positive |
| J07A1120 | Female | 75 | Breast | Invasive ductal carcinoma | Primary | T2 | N1 | M0 | 2B | II | Dead | 59 | 1 | Positive |
| J07A1121 | Female | 33 | Breast | Invasive ductal carcinoma | Primary | T2 | N3 | M0 | 3C | II | Alive | 109 | 1 | Negative |
| J07A1122 | Female | 64 | Breast | Invasive ductal carcinoma | Primary | T2 | N0 | M0 | 2A | II | Alive | 109 | 1 | Negative |
| J07A1124 | Female | 43 | Breast | Invasive ductal carcinoma | Primary | T2 | N2 | M0 | 3A | II | Alive | 109 | 2 | Weakly positive |
| J07A1125 | Female | 71 | Breast | Invasive ductal carcinoma | Primary | T1 | N0 | M0 | 1 | II | Alive | 109 | 1 | Negative |

| | | | | | | | | | | | | | | |
|---|---|---|---|---|---|---|---|---|---|---|---|---|---|---|
| J07A1126 | Female | 49 | Breast | Invasive ductal carcinoma | Primary | T1 | N1 | M0 | 2A | II | Alive | 108 | 1 | Weakly positive |
| J07A1127 | Female | 83 | Breast | Invasive ductal carcinoma | Primary | T2 | N2 | M0 | 3A | II | Alive | 108 | 2 | Negative |
| J07A1128 | Female | 50 | Breast | Invasive ductal carcinoma | Primary | T1 | N0 | M0 | 1 | II | Alive | 108 | 0 | Negative |
| J07A1130 | Female | 54 | Breast | Invasive ductal carcinoma | Primary | T3 | N2 | M0 | 3A | II | Dead | 9 | 3 | Positive |
| J07A1132 | Female | 42 | Breast | Invasive ductal carcinoma | Primary | T1 | N1 | M0 | 2A | II | Alive | 108 | 2 | Positive |
| J07A1133 | Female | 51 | Breast | Invasive ductal carcinoma | Primary | T2 | N0 | M0 | 2A | II | Alive | 107 | 2 | Positive |
